# Phenotype-based screening of synthetic cannabinoids in a Dravet Syndrome zebrafish model

**DOI:** 10.1101/815811

**Authors:** Aliesha Griffin, Mana Anvar, Kyla Hamling, Scott C. Baraban

**Affiliations:** Epilepsy Research Laboratory and Weill Institute for Neuroscience, Department of Neurological Surgery, University of California San Francisco, San Francisco, CA, USA, 94122

## Abstract

Dravet syndrome (DS) is a catastrophic epilepsy of childhood, characterized by cognitive impairment, severe seizures and increased risk for sudden unexplained death in epilepsy (SUDEP). Although refractory to conventional antiepileptic drugs, emerging preclinical and clinical evidence suggests that modulation of the endocanniboid system could be therapeutic in these patients. Here we used a validated zebrafish model of DS, *scn1lab* homozygous mutants, to screen a commercially available library containing 370 synthetic cannabinoid (SC) compounds for compounds effective in reducing spontaneous seizures. Primary phenotype-based screening was performed using a locomotion-based assay in 96-well plates, and a secondary local field potential recording assay was then used to confirm suppression of electrographic epileptiform events. Identified SCs with anti-seizure activity, in both assays, included five SCs structurally classified as indole-based cannabinoids: JWH 018 N-(5-chloropentyl) analog, JWH 018 N-(2-methylbutyl) isomer, 5-fluoro PB-22 5-hydroxyisoquinoline isomer, 5-fluoro ADBICA, and AB-FUBINACA 3-fluorobenzyl isomer. Our approach demonstrates that two-stage phenotype-based screening in a zebrafish model of DS successfully identifies synthetic cannabinoids with anti-seizure activity, and supports further investigation of SCs for refractory epilepsies.

## Introduction

Seizures, in some epilepsy patients, can be difficult to control using conventional antiepileptic drugs (AEDs). In patients classified with catastrophic epilepsies in childhood, effective seizure control using AEDs can be a significant problem and early developmental exposure to AEDs is often associated with undesirable side effects. In the search for new treatment options for these patients there is growing interest in drugs that modulate the endogenous cannabinoid system. The endocannabinoid system, comprised of two G-protein coupled receptors (CB1 and CB2), has been demonstrated to play a role in regulating seizure activity in the brain^1–4^. Recent data suggests broad anticonvulsant activity for synthetic cannabanoids, phytocannabinoids and *Cannabis sativa* (marijiuana) e.g., drugs targeting the endocannabinoid system. Studies on medical marijuana largely focus on anecdotal patient experiences. Careful preclinical examination of medical marijuana and related extracts such as cannabadiol are difficult as these are designated as Schedule II compounds by the U.S. Food and Drug Administration (FDA). Synthetic cannabinoids (SCs), which represent a variety of compounds engineered to bind cannabinoid receptors and minimize the psychogenic effects of marijuana, have been examined most extensively in adult animal models of acute seizures, with mixed results. In the pentylenetetrazole (PTZ) model of generalized seizures, WIN 55,212-2 (a mixed CB1/CB2 receptor agonist), exerts pro- and anticonvulsant effects wherease rachidonyl-2’-chloroethylamide (a CB1 receptor agonist) was shown to decrease acute PTZ seizure thresholds, increase acute PTZ seizure thresholds or have no effect at all^5–8^. In the maximal dentate activation model of limbic seizures or limbic kindling models, WIN 55,212-2 reduced seizure thresholds^1, 2^, delayed epileptogenesis^3^, or failed to provide any seizure protection^4^. Recent testing completed at the Epilepsy Therapy Screening Program reported anti-seizure properties for CBD in mouse 6 Hz 44 mA and mouse/rat maximal electroshock seizure assays^5^. Although even less is known about experimental models of childhood epilepsies, a recent study by^6^ examined seven cannabinoid receptor agonists in young postnatal rat models of PTZ- hypoxia- or methyl-6,7-dimethoxy-4-ethyl-beta-carboline-3-carboxylate(DMCM)-evoked seizures, at concentrations that also induced sedative effects only WIN 55,212-2 exhibited anti-seizure activity in all three models.

Interestingly, Dravet syndrome (DS) - a genetic epilepsy associated, in most cases, with a loss-of-function *de novo* mutation in the *SCN1A* voltage-gated sodium channel subunit - is one particular form of childhood epilepsy where SCs, phytocannabinoids and medical marijuana have consistently shown anticonvulsant activity in preclinical and clinical studies. Reductions in seizure activity were observed in *Scn1a*^+/−^ mice at drug concentrations above 100 mg/kg with amelioration of autistic-like behaviors seen at a lower concentration of 10 mg/kg^7^. In the first open-label investigational trial of cannabidiol (CBD) in children with DS, Devinsky et al.^8^ reported a 37% median reduction in monthly seizure counts. Additional positive seizure reductions with CBD treatment^9–11^, and a GW Pharmaceuticals sponsored open-label double-blinded study reporting an adjusted 23% reduced in seizure frequency led to FDA approval for CBD (Epidiolex^®^) as a treatment for seizures associated with DS.

In the present study, we used a *scn1* zebrafish mutant model of DS to screen a library of SCs for those that exhibit anti-seizure properties. The *scn1lab* mutant zebrafish line exhibits spontaneous seizures that can be easily monitored using acute behavioral and electrophysiological assays^12, 13^. These mutants exhibit metabolic deficit, early fatality, sleep disturbances and a pharmacological profile similar to DS patients^14–16^. In addition to replicating key aspects of DS, *scn1lab* mutants were shown to be a useful model system for large-scale phenotype-based drug screening and drugs identified with this zebrafish-centric approach have already shown efficacy in the clinic^13, 17^. Using *scn1lab* mutants, we screened 370 SCs at two different concentrations. After repeated locomotion and electrophysiological assays with multiple biological replicates, we identified five SCs that exert significant anti-seizure activity (reducing convulsive swim behavior and suppressing abnormal electrographic discharge frequency) during acute exposures. Additionally, analysis of structure-activity relationships revealed these compounds are structurally similar and are indole-derived cannabinoids^18^. These data suggest that specific classes of SCs can exert antiepileptic activity and provide further justification for using larval zebrafish models to identify novel therapeutic targets.

## Materials and Methods

### Zebrafish maintenance

Zebrafish were maintained according to standard procedures^19^ and following guidelines approved by the University of California, San Francisco Institutional Animal Care and Use Committee. The zebrafish room was maintained on a light-dark cycle, with lights-on at 9:00 AM and lights-off at 11:00 PM. An automated feedback control unit was used to maintain aquarium water conditions in the following ranges: 29-30°C, pH 7.5-8.0, conductivity (EC) 690-710. Zebrafish embryos and larvae were raised in an incubator maintained at 28.5°C, on the same light-dark cycle as the fish facility. Water used for embryos and larvae was made by adding 0.03% Instant Ocean and 0.000002% methylene blue to reverse-osmosis distilled water. Embryos and larvae were raised in plastic petri dishes (90 mm diameter, 20 mm depth) and housing density was limited to approximately 50-60 individuals per dish. The sex of embryos and larvae cannot be determined at these early stages.

### Drugs

Compounds for drug screening were purchased from Cayman Chemicals and were provided as 10 mM DMSO solutions (Supplemental Table I).

### Locomotion assay

At 5 days post fertilization (dpf) offspring from crossing adult *scn1lab*^+/−^ zebrafish were sorted by pigmentation to isolate *scn1lab*^−/−^ larvae^15^ and placed individually in one well of a 96-well plate (75 μL of E3 media). The plate was positioned in a DanioVision (Noldus) chamber under dark light for a 20-min habituation period before obtaining a 10 min “Baseline Recording” epoch. Fresh drug solutions were prepared on each day of experimentation in 1 mL of E3 media. After baseline measurements, embryo media was carefully removed and 75 μL test drug (at a concentration of 10 or 250 μM) or embryo media (control) were added. The plate was returned to the chamber for another 20-min habituation period before beginning a 10 min “Experimental Recording” epoch. Criteria for a positive hit designation were as follows: (1) a decrease in mean velocity of ≥40%, and (2) a reduction to Stage 0 or Stage I seizure behavior (defined in Baraban et al. 2005^28^) in the locomotion plot for at least 50% of the test fish. Each test compound classified as a “positive hit” in the locomotion assay was assessed for toxicity by direct visualization on a stereomicroscope following a 60 min drug exposure. Acute toxicity (or mortality) was defined as no visible heartbeat or movement in response to external stimulation in at least 50% of the test fish. Compounds identified as successful in the first locomotion screen were tested on an independent clutch of larvae using the method described above. Compounds that were successful in two independent locomotion assays, and were not acutely toxic, were tested a third time using fresh drug stock sourced from Cayman Chemical.

### Electrophysiology assay

Zebrafish larvae were first monitored in the 96-well format locomotion assay described in 2.3. Individual larvae were then briefly exposed to cold and immobilized in 1.2% agarose by an investigator blinded to the locomotion assay parameter. Local field potential recordings (LFP) were obtained from midbrain using a single-electrode recording, as previously described^20, 21^. LFP recordings, 10 min in duration, were obtained using Axoclamp software (Molecular Devices; Sunnyvale, CA) at an acquisition rate of 1 kHz. Abnormal electrographic seizure-like events were analyzed *post hoc* as: (i) brief *interictal-like* events (0.47 ± 0.02 sec duration; n = 52) comprised of spike upward or downward membrane deflections greater than 3× baseline noise level or (ii) long duration, large amplitude *ictal-like* multi or poly-spike events (3.09 ± 1.01 sec duration; n = 21) greater than 5× baseline noise level. Noise level was measured as 0.35 ± 0.03 mV (n = 18). All *scn1lab* mutants exhibit some form of spontaneous electrographic seizure activity and no clustering of activity was observed even with prolonged electrophysiology monitoring (see^22^). Both events were counted using “threshold” and/or “template” detection settings in Clampfit (Molecular Devices; Sunnyvale, CA). All embedded larvae were continuously monitored for blood flow and heart rate using an Axiocam digital camera. All experiments were performed on at least two independent clutches of *scn1lab* zebrafish larvae (5-6 dpf).

### Statistics

Electrophysiological data was examined by one-way analysis of variance with subsequent Dunnett multiple comparisons tests.

## Results

### Behavioral screening

To identify SC compounds that modify convulsive swim behavior, we performed an *in vivo* primary screen using an automated locomotion tracking protocol^12, 13^. Mutants were placed individually in a single well of a 96-well plate and a baseline locomotion tracking plot (10 min in duration) was obtained in embryo media. Embryo media was removed and mutants were then exposed to test SC compounds at a single screening concentration of 250 μM (n = 6 fish/drug). All compounds remained coded by compound number provided by Cayman Chemical. Mutant swim activity between two consecutive recording epochs in embryo media was tracked on every plate as an internal control. Mean velocity was calculated for each well and the percent change from baseline for all 370 test compounds is plotted in Figure 1a; drugs that were toxic at 60 min are shown in red. On first pass locomotion screening at 250 μM, 27% of SC compounds were identified as positive hits and 39% of SC compounds were identified as acutely toxic (Figure 1c). To reduce the percent of compounds being identified as toxic and potential false negatives, a second screen was perfomed on all 370 compounds at a concentration of 10 μM (n = 6 fish/drug), following the same protocol (Figure 1b). On first pass locomotion screening at 10 μM, 25% of SC compounds were identified as acutely toxic and 22% were identified as positive hits (Figure 1c). There were 20 SC compounds which effectively decreased the high velocity seizure-like swim behavior in both the 10 μM and 250 μM behavior screen (Fig 1d,e).

**Figure 1.**
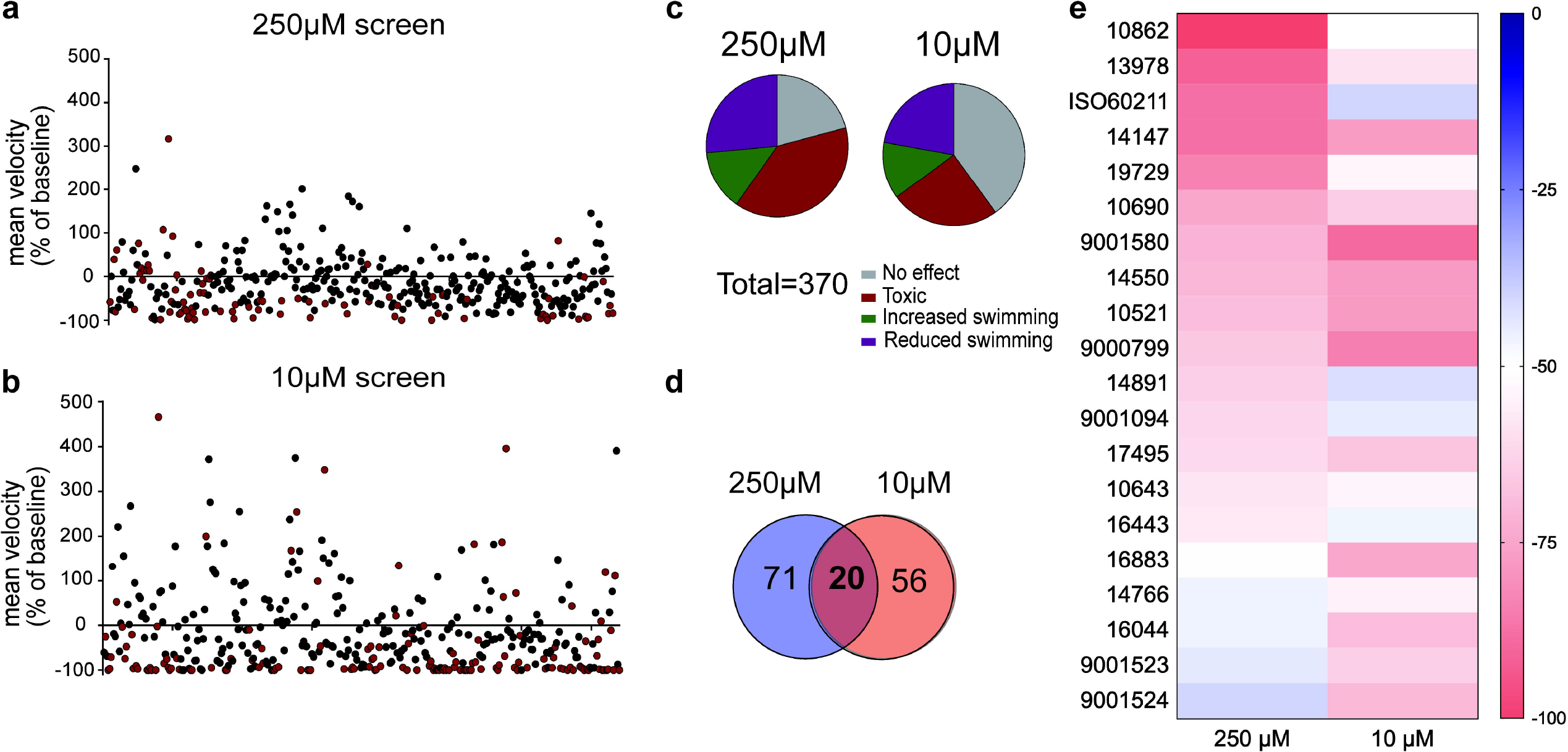
A library of synthetic cannabinoids was screened for their ability to reduce the high velocity seizure-like swim behavior of 5 day old zebrafish larvae. Compounds were screened at (a) 250 μM, and (b) 10 μM. Each data point represents the mean velocity change in swim behavior of six fish treated with an individual compound. The red data points represent compounds that failed to go into solution or identified as toxic after 90-min exposure. (c) Summary of compound effects after screening 370 synthetic cannabinoids at 250 μM and 10 μM. The threshold for inhibition of seizure activity was determined as a reduction in mean swim velocity of ≥40%. (d) The total number of compounds identitied as positive from the 250 μM and 10 μM library screen. (e) Heat map representing the 20 compounds which successfully reduced the mean swim velocity by >40% in the 250 μM and 10 μM screening.

Next, all 20 SCs compounds which emerged from the two primary screens were uncoded and sourced as individual drugs from Cayman Chemical for a concentration-response studies using *scn1lab* mutant larvae. SCs were tested at 1, 10 and 100 μM (n = 6 fish/drug/concentration). A threshold for positive hits was set as a decrease in mean velocity of 40% or more. Additionally, a Stage 0 or I swim behavior and a dose response needed to be observed. Representative locomotion tracking plots are shown in Figure 2b. Five SC compounds were identified as being able to decrease the seizure-like swim behavior observed in the *scn1lab* mutant larvae in a dose dependent manner (Figure 2). Aggregating data from the first stage of our zebrafish antiseizure drug screening platform, these five SC compounds were classified as positive hits for reducing seizure-like swim behavior and subsequently moved on to the electrophysiology assay for confirmation.

**Figure 2.**
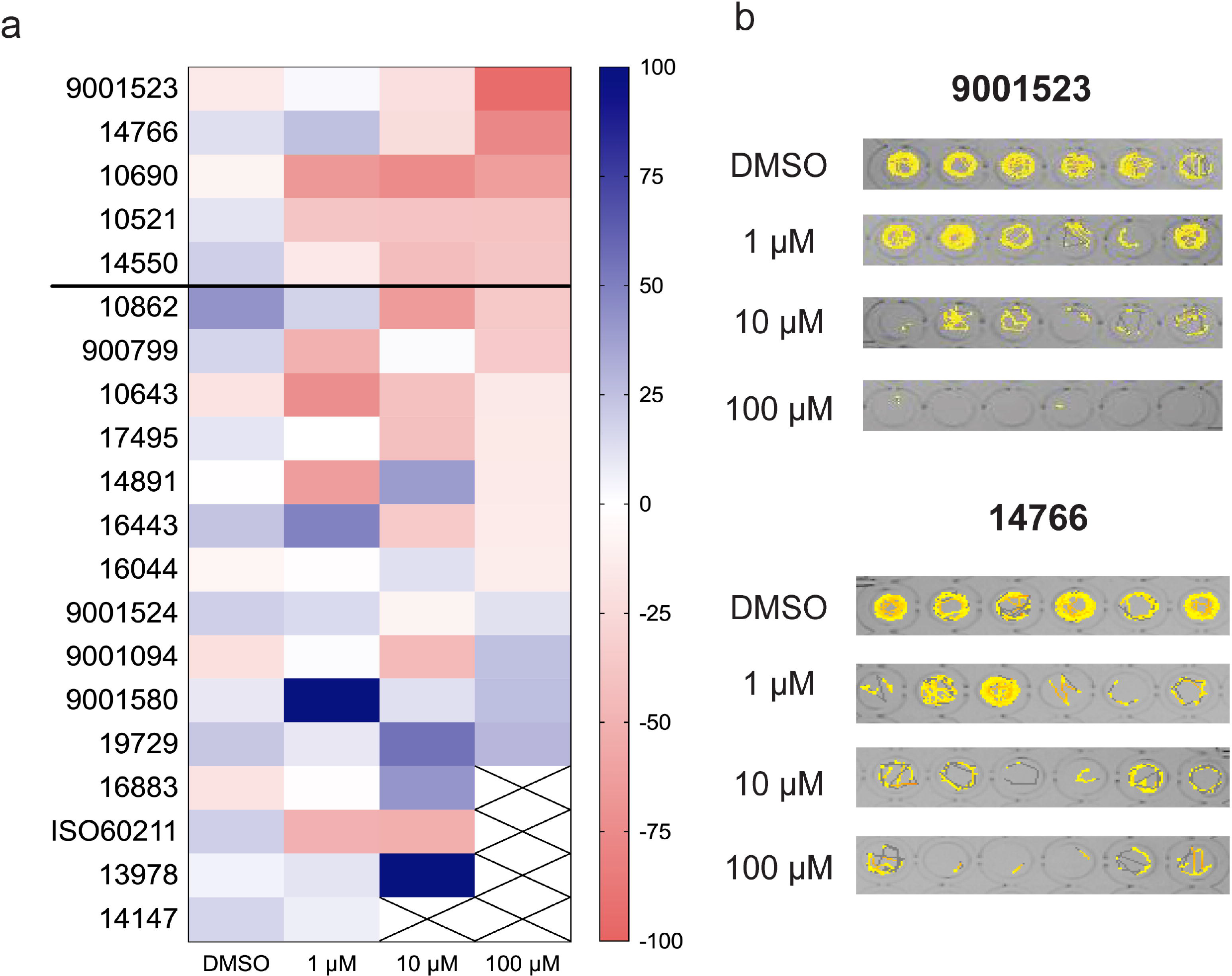
Behavioral screening of 20 compounds which were identified as positive from the library screens. (a) Heat map representing the change of mean swim velocity of the 20 hit compounds. The 20 synthetic cannabinoids which were identified from the blind screens were retested at 1μM, 10μM and 100μM to confirm a dose response effect. Compounds marked with a cross were identified as toxic. (b) Representative swim behavior traces obtained during a 10 min recording epoch for the top two compounds, #9001523 and #147666.

### Electrophysiology screening

To evaluate whether selected SCs inhibit seizures, we performed an *in vivo* secondary screen using electrophysiology techniques^12, 13, 23^. Mutants were placed in a single well of a 96-well plate and exposed to test SC compounds (#10521, #10690, #14550, #14766, #900799 and #9001523; Table 2) at drug concentrations determined above or DMSO (vehicle). Compound #900799 was selected here as a “control” SC that fell below the positive hit threshold described above for the final locomotion-based assay. After a 30 min exposure, *scn1lab* mutant larvae were immobilized in agar for LFP recording and *post hoc* analysis of both interictal- (range: 107-453 events/10 min) and ictal-like (range: 1-10 events/10 min) activity. Exposure to compounds #10521, #10690, #14500, #14766 and #9001523 significantly reduced the frequency of these spontaneous epileptiform events (*F*_(6,80)_ = 14.41, ***p < 0.0001 or **p < 0.0003) (Figure 3a). Representative 10 min recording epochs for each compound, and an age-matched *scn1lab* mutant larvae control are shown in Figure 3B. Note the presence of robust interictal- and ictal-like events in *scn1lab* controls and following exposure to compound #900799 but the near complete absence of events with exposure to five different SCs.

**Figure 3.**
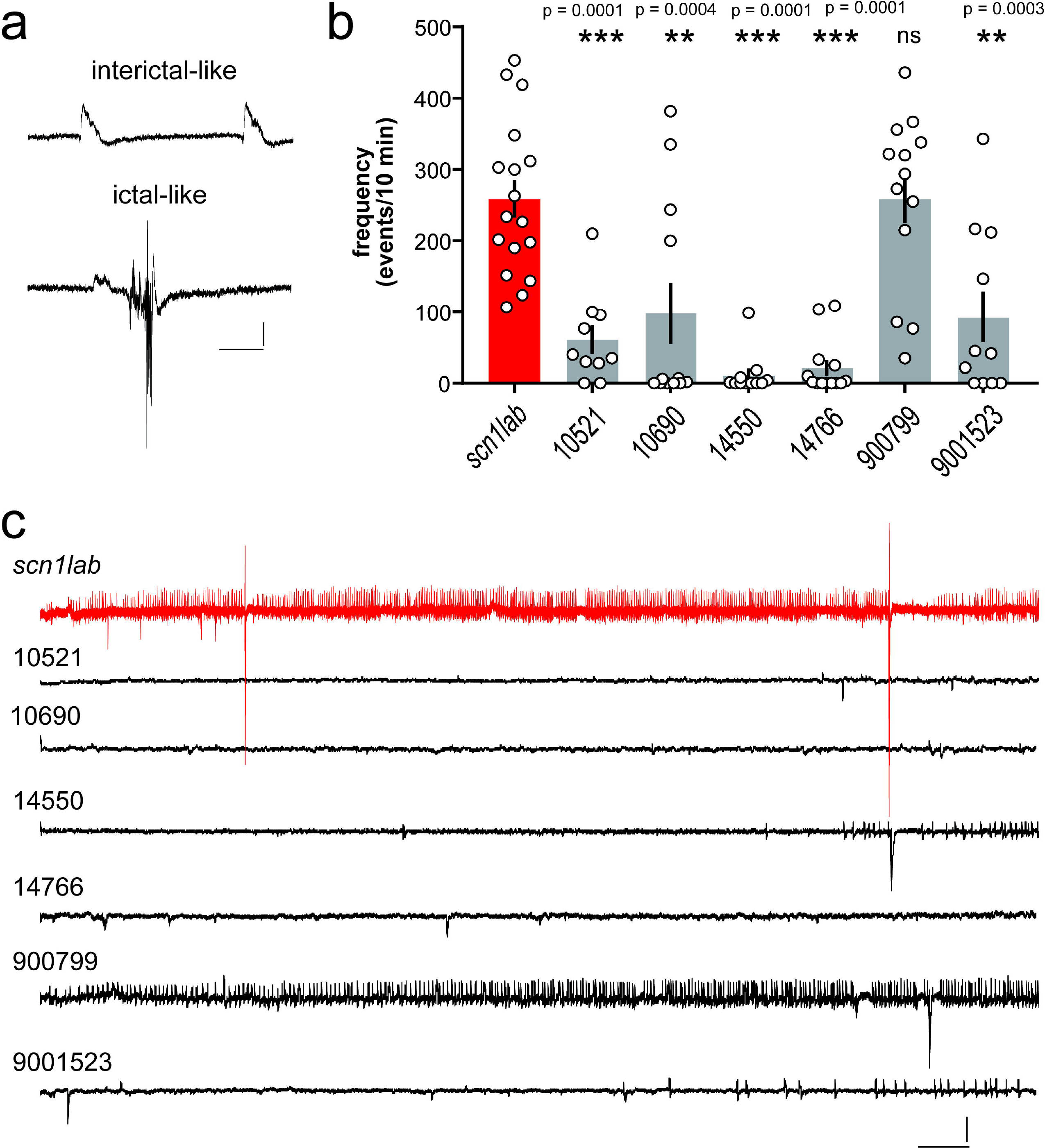
Electrophysiological assay for compounds identified in the locomotion-based screening assay. (a) Local field potential (LFP) recordings were obtained with an glass micro-electrode placed under visual guidance in the midbrain of agar-immobilized *scn1lab* larvae that had previously showed reduced seizure-like behavior in the locomotion assay. Representative examples of events classified as interictal- or ictal-like are shown. Scale bars: 1 mV, 0.5 sec. (b) Bar graphs show the frequency of epileptiform events in a 10 min recording epoch for *scn1lab* larvae exposed to DMSO vehicle (scn1lab mutants; n = 17), JWH 018 N-(5-chloropentyl) analog (compound #10521; n = 10), JWH 018 N-(2-methylbutyl) isomer (compound #10690; n = 12), 5-fluoro PB-22 5-hydroxyisoquinoline isomer (compound #14550; n = 11), 5-fluoro ADBICA (compound #14766; n = 13), JWH 018 adamantyl analog (compound #9000799; n = 13), and AB-FUBINACA 3-fluorobenzyl isomer (compound #90001523; n = 11). Mean ± SEM and individual data points are shown. One-way analysis of variance with Dunnet’s multiple comparisons was used to test for significance. **p < 0.001; *** p < 0.0001. (c) Representative electrophysiology traces (10 min) are shown for SC compouns 10521, 10690, 14550, 14766, 9000799 and 900015323 compared to an *scn1lab* mutant zebrafish (red). Scale bars: 1 mV, 10 sec.

**Figure 4.**
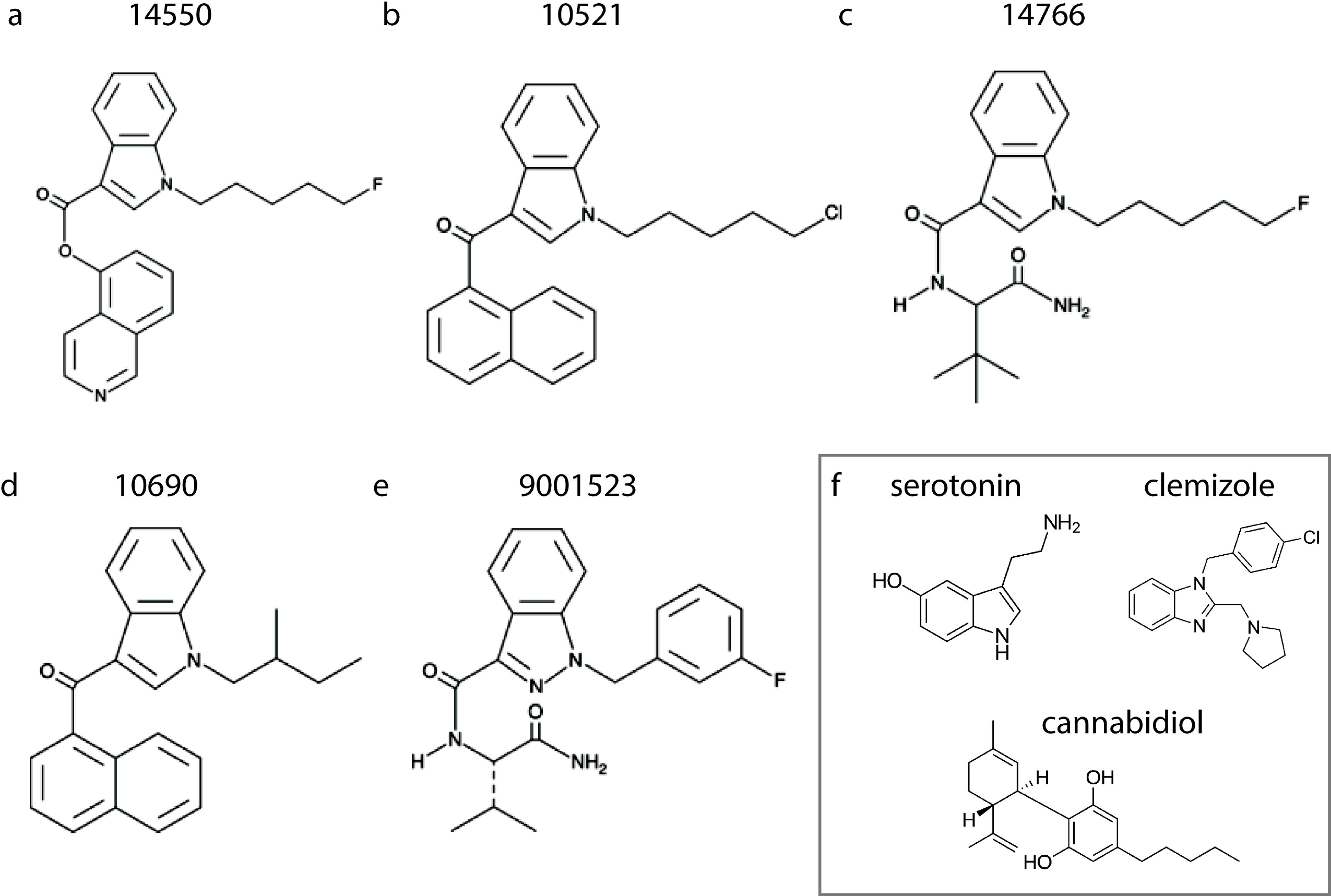
Structural comparison of SC identified to reduce spontaneous seizures in the *scn1lab* Dravet syndrome zebrafish mutant. (a) 14550; 5-fluoro PB-22 5-hydroxyisoquinoline isomer, (**b**) 10521; JWH 018 N-(5-chloropentyl) analog, (**c**) 14766; 5-fluoro ADBICA, (**d**) 10690; JWH 018 N-(2-methylbutyl) isomer, and (**e**) 9001523; AB-FUBINACA 3-fluorobenzyl isomer. (f) These indol derived cannabinoids share structural similarity with serotonin and clemizole (currently in clinical trial for Dravet syndrome). They are structurally unrelated to cannabidiol.

**Table 1:** Synthetic cannabinoid compounds identified in a two-stage screening process

| Compound No. | Compound Name | Description |
| --- | --- | --- |
| 10521 | JWH 018 N-(5-chloropentyl) analog | JWH 018 is a synthetic cannabinoid that potently activates the central cannabinoid (CB1) and peripheral cannabinoid (CB2) receptors ($K_i = 9.0$ and $2.94$ nM, respectively). JWH 018 N-(5-chloropentyl) analog differs structurally from JWH 018 by having chlorine added to the 5 position of the pentyl chain. |
| 10690 | JWH 018 N-(2-methylbutyl) isomer | JWH 018 2-methylbutyl homolog is an analog of JWH 018, a mildly selective agonist of the central cannabinoid receptor ($K_i = 8.9$ nM) derived from the aminoalkylindole WIN 55,212-2. |
| 14550 | 5-fluoro PB-22 5-hydroxyisoquinoline isomer | 5-fluoro PB-22 is an analog of 5-fluoropentyl JWH-type cannabimimetics. 5-fluoro PB-22 5-hydroxyisoquinoline isomer differs from 5-fluoro PB-22 by having the quinoline group replaced with an isoquinoline group attached at its 5 position. |
| 14766 | 5-fluoro ADBICA | This compound is a derivative of ADBICA featuring a fluorine atom added to the terminal carbon of the pentyl group. |
| 9000799 | JWH 018 adamantyl analog | JWH 018 adamantyl analog is a mildly selective agonist of the peripheral cannabinoid receptor, where the naphthalene ring is substituted with an adamantyl group. |
| 9001523 | AB-FUBINACA 3-fluorobenzyl isomer | AB-FUBINACA is an indazole-based synthetic cannabinoid that potently binds the central CB1 receptor ($K_i = 0.9$ nM). |

## Discussion

Here we describe the first large-scale phenotype-based screen of a synthetic cannabinoid library using a zebrafish model of Dravet syndrome. Our approach to screening compounds for anti-seizure activity in *scn1lab* mutant zebrafish builds on earlier model validation^12^, acute drug exposure protocols^17^, and a database now exceeding 3200 compounds^13, 17, 23, 24^. Interest in cannabinoid-based treatment options for DS began with anecdotal reports and ultimately led to FDA approval of cannabidiol (Epidiolex^®^) for the treatment of seizures associated with this disorder. Using a stringent two-stage screening strategy, we identified five SCs that exert significant anti-seizure activity in *scn1lab* mutant zebrafish. SCs add to a growing list of DS relevant drugs successfully identified in *scn1* zebrafish that include “standard-of-care” AEDs (e.g., valproate, stiripentol, benzodiazepines, bromides) and experimental drugs (e.g., fenfluramine, lorcaserin, trazodone, clemizole).

Synthetic cannabinoids (SCs), first developed in the 1940s, represent a variety of compounds engineered to bind cannabinoid receptors. SCs bind G-protein coupled CB1 receptors located at the presynaptic terminal and are also thought to interact with voltage-gated potassium channels, voltage-gate sodium channels, N-and P/Q-type-calcium channels, and an orphan G-protein coupled receptor (GPR55). Whether one, or several, of these potential sites of action are responsible for the anti-seizure effects noted with SCs (or CBD) remains a controversial area of investigation. *In vivo* studies using intracerebroventricular administration of a CB1-receptor agonist, arachidonyl-2-chloroethylamide (ACEA), significantly decreased the frequency of penicillin-induced epileptiform activity in rats^25^. CMYL-4CN-BINACA, another CB1 receptor agonist, elicited pro-convulsant effects in mice^26^ and a synthetic cannabinoid (AM2201) induced epileptiform activity and convulsive behaviors in mice that could be blocked by the selective CB1 receptor antagonist AM251^27^. Investigating alternatives to a CB1/2 receptor mechanism, Kaplan et al.^7^ showed that antagonism of GPR55 receptors mimic the effects of CBD *in vitro* and could explain anti-seizure effects of high concentrations of CBD in *Scn1*^+/−^ mice. However, Huizenga et al.^6^ reported that nonselective CB1/2 and selective CB1 agonists suppress activity in neonatal seizure models (though these findings were not replicated with GPR55 agonists). In patch-clamp recordings from isolated mouse cortical neurons, CBD decreased neuronal firing activity and significantly reduced peak whole-cell voltage-activated sodium currents^28^. In contrast, in a recent study using rodent and human induced pluripotent stem cell (iPSC)-derived excitatory or inhibitory neurons, CBD reduced neuronal firing activity but did not alter voltage-gated sodium channel conductance^29^.

Zebrafish possess orthologues for 82% of human disease-associated genes^30^. Orthologous zebrafish proteins are often similar to human within their functional domains. For example, between 85% and 95% of the amino acids in a ligand-binding domain of the zebrafish glucocorticoid receptor are similar to those in human. These properties combined with the ease of large-scale pharmacological screening make larval zebrafish a good model system for investigating drug targets. A recent study^31^, limited only to locomotion assay read-outs in larval zebrafish, reported a synergistic effect of Δ-9-tetrahydrocannabinol (THC) and cannabidiol (CBD) on hyperactive behaviors. Unfortunately, these studies failed to include electrophysiology measures of efficacy and were limited to only these two compounds. Here we screened 370 different synthetic cannabinoids in a locomotion assay at two different drug concentrations, initially identifying 20 compounds as exerting potential therapeutic benefit. With repeated biological replicates, newly sourced compound re-tests and sensitive secondary electrophysiologay assays, we narrowed this initial list to five SCs that were classified as effective in suppressing spontaneous seizures (behavioral and electrographic) in *scn1lab* mutants. This scientifically stringent two-stage approach further validates the selectivity of our screening strategy as only 1.3% of all SCs tested were identified as positive hits. A caveat of this strategy is that pharmacokinetic data for how these SCs are absorbed, distributed or metabolized in larval zebrafish is not available which would suggest an error in false negative designations. Interestingly, as several of these SCs are classified as indole-derivatives these data also suggest a potential target not recognized in previous preclinical studies. Indole-derived cannabinoids have strong binding affinity for the 5HT_2B_ receptor^32^. This serotonin receptor subtype was recently identified by our group as potentially mediating the anti-seizure effects of clemizole^33^. Taken together, a convergence of evidence now exists to suggest that activation of a serotonin 2B receptor is a potential target for DS therapy.

Although further preclinical studies investigating SCs identified here are warranted, a note of caution needs to be extended to clinical translation of these data. Substantial evidence of serious adverse effects accompany these compounds. These include, and may not be limited to, panic attacks, memory distortions, paranoia, psychotic reactions, disorganized behavior, and suicidal thoughts^34, 35^. Reports of acute ischemic stroke^36, 37^, seizures^38, 39^, and sudden death^40–42^ with SCs are not uncommon. In particular, indole- and indazole-based SCs are associated with clinical signs of reduced consciousness, paranoia and seizures in humans^43^. Acknowledging these limitations, the current studies provide tools for further preclinical research investigating the anti-seizure potential of the endocannabinoid signaling system. This information is valuable for investigators and clinician-scientists interested in the potential benefits of cannabinoid receptor agonists for intractable seizure disorders.

## Acknowledgements

This work was supported from NINDS R01 grants no. NS079214 and NS103139 (S.C.B). We would like to thank members of the Baraban laboratory for their useful discussions during the course of these studies; Matthew Dinday, Chinwendu Ononuju and Harrison Ramsay in particular for their support in zebrafish facility maintenance.

## Author contributions

Locomotion-based screening assays were performed by M.A., K.H. and A.G. Electrophysiology experiments were performed by S.C.B. The design, conceptualization, interpretation of data, data analysis and writing of the manuscript was done by A.G. and S.C.B. All authors contributed to the editing process.

## Data availability

The datasets generated and analyzed in the current study are available from the corresponding author on reasonable request.

